# Endogenous expression of mutant RagC in mice results in early-onset dilated cardiomyopathy with features of the Long-Olsen-Distelmaier syndrome

**DOI:** 10.64898/2026.11.09.765270

**Authors:** Ana Belén Plata-Gómez, Elena Sánchez García, Ana Sagrera, Lucía de Prado-Rivas, Ayelén M. Santamans, Alba Sanz, Ana Ortega-Molina, Lucas Fellmann, Alba C. Arcones, Luis Leiva-Vega, Eduardo Caleiras, Elena Piñeiro-Yáñez, Sara Cogliati, Guadalupe Sabio, Alejo Efeyan

## Abstract

Dilated cardiomyopathy (DCM) is a severe heart condition often linked to arrhythmias, clots, and heart failure, for which treatments are not yet curative. Recently, *de novo* mutations in genes encoding components of the Rag GTPase nutrient signaling pathway, particularly in *RRAGC*, have been found in unrelated patients with a severe, early-onset DCM syndrome called Long-Olsen-Distelmaier. Cancer cell lines, hiPSC-derived cardiomyocytes and zebrafish have been used to study this *Ragopathy*, but a mammalian model to investigate this syndrome has not been generated. Here, we show that mice engineered to endogenously express mutations in Serine 74 (orthologous to the Serine 75 mutated in patients), exhibit early-onset features of DCM with decreased heart size, fibrosis, diastolic dysfunction and aortic insufficiency, characteristics reminiscent of the human syndrome. Moreover, *Rragc*-mutant hearts display decreased left ventricular mass and interventricular septum. Endogenous heterozygous expression of *Rragc* mutant variants result in a milder, sublethal defective physiological cardiac growth with increased systolic and diastolic blood pressure. Mechanistically, active RagC does not cause abnormal increase in cardiomyocyte size, but leads to suppressed transcriptional activity of Tfeb and alterations in cardiomyocyte metabolism. Transcriptional profiling of neonatal mutant hearts reveals multiple signatures of impaired mitochondrial activity. *Ex vivo* and *in vitro* analyses of murine cardiomyocyte mitochondria from *Rragc*-mutant mice show suppressed mitochondrial respiration, disturbed complex IV multimerization and super-complex organization. These data point to abnormal early postnatal heart metabolism by deregulated nutrient – Rag GTPase signaling with a potential molecular explanation to the early onset of DCM in patients with *de novo* missense mutations in *RRAGC*, and suggests the relevance of early metabolic interventions in conditions with altered nutrient signaling.

## Introduction

Dilated cardiomyopathy (DCM) is a group of diseases characterized by impaired heart contractility, left ventricular dilation without abnormal loading or coronary disease^1,2^. DCM is a main cause of morbidity and mortality, as it has a strong association with heart failure and lethal arrythmias, and is also a leading cause of heart transplantation^3,4^. DCM is the most common form of pediatric cardiomyopathies^5^, and monogenic and polygenetic drivers have been proposed to account to for around 40% of DCM^6–8^. These drivers include both germline and *de novo* mutations in genes important for structural and functional cardiac homeostasis^3,9^. Recently, several unrelated cases of a DCM syndrome in patients carrying *de novo* single-nucleotide, missense mutations in the gene *RRAGC* have been reported^10,11^, and this syndromic disease has been recently coined Long-Olsen-Distelmaier syndrome (OMIM #620609) (https://omim.org/entry/620609). *RRAGC* encodes for the RagC GTPase, an obligate heterodimeric partner of RagA and a molecular switch that connects cellular nutrient sufficiency to anabolic growth by activating the mechanistic target of rapamycin complex 1 (mTORC1)^12,13^. When nutrients (several amino acids, glycolytic intermediates and lipids) are plentiful, convergent signal transduction cascades result in the GTP loading of RagA and the GDP loading of RagC^14–16^. This nucleotide configuration results in the binding of the Rag GTPase complex to mTORC1 and its recruitment to the outer lysosomal surface, an essential step required for mTORC1 activation^17^. Interestingly, the *de novo* mutations identified in Long-Olsen-Distelmaier syndrome are all missense and overlap with a spectrum of point mutations in *RRAGC* sporadically mutated in human cancer, which functionally mimic a GDP-bound, active state of RagC, and result in a moderate activation of mTORC1^18–20^. The effect of Rag GTPase mutations in cancer cells and in cardiomyocytes, have been interrogated using cultured cells, human induced pluripotent stem cells (hiPSC), and a zebrafish system^10,11,18,19,21–24^. Thus, previous research has illustrated the gain-of-function nature of these mutations, and provided valuable insight of molecular mechanisms downstream of anomalous function of RagC^25^, but a mammalian *in vivo* model of the human condition has not been generated.

We have previously engineered the mouse genome to knock-in gain-of-function mutations of *Rragc* with endogenous regulation. Heterozygous expression of *Rragc* mutants result in a modest but consistent activation of mTORC1 that fosters tumor development^24^, and also contributes to parenchymal damage across organs and results in shortened longevity^26^. Among the mutations engineered in the mouse genome, serine 74 and threonine 89 are orthologous to serine 75 and threonine 90 mutated in Long-Olsen-Distelmaier syndrome, so we hypothesized that heterozygous *Rragc* mutant (*Rragc^mut/+^*) mice may sub-lethally recapitulate aspects of DCM. Indeed, we found that *Rragc*-mutant mice exhibit early-onset DCM, with abnormal heart structure and cardiac damage, thus partially mirroring the human condition. Molecular analysis of *Rragc^mut/+^*hearts underscores abnormal mitochondrial function and respiration complex assembly. Thus, we provide here a mouse model with extensive features of Long-Olsen-Distelmaier syndrome, demonstrate the causative role of activating mutations in *RRAGC* and its consequences for mitochondrial and cardiomyocyte homeostasis.

## Results

### Mutations in the RagC GTPase in mice drive early-onset DCM

The independent identification of *de novo*, somatic mutations in early-onset DCM patients strongly suggests that abnormally increased nutrient signaling during heart development or maturation may be a driver of heart disease. Thus, we took advantage of two strains of knock-in mice harboring heterozygous mutation in amino acidic position 74, namely S74C and S74N, orthologous to the human S75 mutations found in two unrelated patients with Long-Olsen-Distelmaier syndrome with infantile fatal DCM^10,11^. Comprehensive phenotypic and mechanistic analyses were performed primarily in *Rragc^S74N/+^*mice. In addition, key findings were additionally validated in the *Rragc^S74C/+^*line, while ensuring efficient use of animals in accordance with the 3R principles (Replacement, Reduction, and Refinement). *Rragc^mut^* mice were previously shown to present decreased survival^10,11^. Intriguingly, *Rragc^mut^* mice are found at sub-mendelian ratios^24,26,27^ at weaning, but proportions were normal when genotyped at birth (Figure S1a and 1b), as reported previously for other gain-of-function alleles in the Rag GTPase pathway. While this partially-penetrant lethality does not provide specific support for our hypothesis, it is consistent with the early onset of the reported clinical cases and is an expected outcome for a potential underlying early onset heart disease in *Rragc^mut^* mice. To determine whether mTORC1 activity was increased in hearts from *Rragc^mut^* mice, we first quantified the phosphorylation of its downstream target S6 in whole protein extracts from *Rragc^+/+^*and *Rragc^S74N/+^* hearts. As previously reported for organs and primary cells derived from *Rragc^mut/+^* mice^24,26^, phospho-S6 was significantly increased upon fasting in *Rragc^S74N/+^*samples, as compared to wild-type littermates (Figure S1c). MTORC1 activation is associated with increase in cell size across tissues^28–30^ but cardiomyocyte-specific deletion of RagA and RagB results in increased heart weight^31^. Thus, we next measured heart weight in neonates (E19.5) and in adult (8-15 week-old) mice, and instead of observing mTORC1-induced organ hypertrophy, *Rragc*-mutant hearts were significantly smaller (Figure 1a and 1b, and Figure S1d and 1e). We hypothesized that decreased heart weight could be either a consequence of abnormal heart development or associated with abnormal cardiomyocyte size, but quantification of cardiomyocyte cross sectional area revealed non-significant changes in cardiomyocytes size in *Rragc*-mutant hearts (Figure 1c and 1d, and Figure S1f and S1g). Instead, and consistently with a moderate cardiac damage, a significant increase in fibrosis was evident in *Rragc^S74N/+^* young hearts (Figure 1e). To find direct evidence for an underling, DCM-causing abnormal structure and function of the heart, we performed repeated echocardiographic analysis in mice from 4 to 15 weeks old (Figure 1f). Strikingly, *Rragc^S74N/+^* mice showed diastolic dysfunction (Figure 1g and Figure S1h) and a degree of aortic insufficiency (Figure 1h and Figure S1i). Left ventricular mass was decreased (Figure 1i), consistently with the decreased heart weight observed in Figure 1a and 1b. Another key structural feature of DCM is the thinning of the left ventricular walls, and accordingly, the interventricular septum thickness (IVS,d) and the left ventricular posterior wall (LVPW,d), (Figure 1j and 1k) were decreased in *Rragc*-mutant hearts, without compromised left ventricular internal diameter end diastole (LVID,d; Figure S1j) and without decreased ejection fraction (EF, Figure S1k). The LVID,d/LV mass ratio was increased (Figure 1l and Figure S1l), indicating greater ventricular dilation relative to wall thickening. Finally, blood pressure was found increased in adult *Rragc^S74N/+^*mice (Figure 1m).

**Figure 1.**
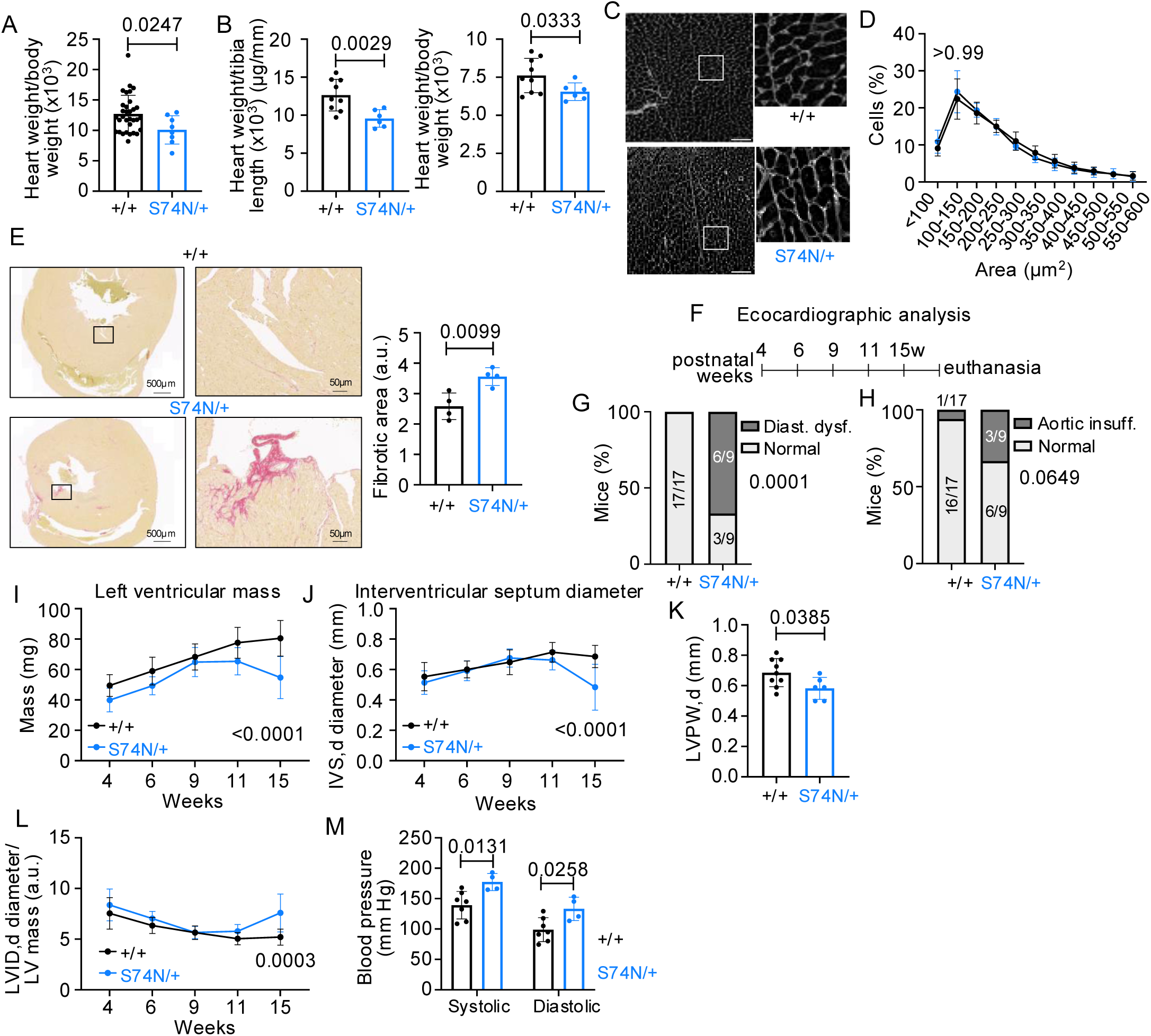
Activation of the nutrient signaling pathway in *RragcS74N/+* mice drives an early-onset of DCM. **A.** Heart weight/body weight ratio of E19.5 *Rragc+/+* (n=32) and *RragcS74N/+* (n=8) neonates. **B.** Heart weight/tibia length (left) and heart weight/body weight (right) ratios of 15 week-old *Rragc+/+* (n=9) and *RragcS74N/+* (n=6) mice. **C.** Cardiomyocyte size in 15 week-old mice analyzed by WGA staining. Scale bar 75μm. **D.** Quantification of cardiomyocyte size in *Rragc+/+* (n=4) and *RragcS74N/+* (n=4) mice from C. **E.** Representative picrosirius red staining in heart slides from 15 week-old *Rragc+/+* and *RragcS74N/+* mice (left). Quantification of fibrosis in *Rragc+/+* (n=4) and *RragcS74N/+* (n=4) (right). **F.** Experimental design for echocardiographic analysis of *Rragc+/+* (n=17), *Rragc^S74N/+^* (n=9) mice. **G.** Diastolic dysfunction found by echocardiographic analysis from 4 to 15 week-old in *Rragc^+/+^* (n=17) and *RragcS74N/+* (n=9) mice. **H.** Aortic insufficiency found by echocardiographic analysis from 4 to 15 week-old in *Rragc+/+* (n=17) and *RragcS74N/+* (n=9) mice. **I.** Left ventricular mass measured by echocardiographic analysis from 4 to 15 week-old in *Rragc+/+* (n=9) and *RragcS74N/+* (n=6) mice. **J.** Interventricular septum diameter measured by echocardiographic analysis from 4 to 15 week-old in *Rragc+/+* (n=9) and *RragcS74N/+* (n=6) mice. **K.** Left ventricular posterior wall measured by echocardiographic analysis from 15 week-old in *Rragc+/+* (n=9) and *RragcS74N/+* (n=6) mice. **L.** LVID,d/LV mass ratio measured by echocardiographic analysis in *Rragc+/+* (n=9) and *RragcS74N/+* (n=6) mice. **M.** Blood pressure in 15 week-old *Rragc+/+* (n=7) and *RragcS74N/+* (n=4) mice. In all panels data are presented as mean values ± SD. Statistical significance was calculated by using unpaired two-tailed *t*-test in A, B, C, E and K, 2-way ANOVA with Sidak’s multiple comparisons test in D, I, J, L and M and chi-squared in G and H. p values are indicated in each panel.

Altogether, these results support that activating mutations in *Rragc* orthologous to those derived from early-onset DCM seen in Long-Olsen-Distelmaier syndrome result in abnormal heart development and in compromised cardiac homeostasis. The observation of cardiac damage with an increase in the deposition of collagen fibers in young animals is likely to compromise the relaxation capacity of the left ventricle, resulting in altered ventricular pressure. In turn, elevated pressure drives aortic insufficiency and diastolic dysfunction. These pathologies are likely responsible for the observed structural alterations in the heart, specifically leading to a decrease in left ventricular mass and wall thickness, giving rise to an early dilated cardiomyopathy-like phenotype similar to that observed in patients of this Ragopathy.

### Effect of *Rragc* mutations in old mice

To determine the evolution of the consequences of increased nutrient signaling in the mouse heart, we assessed structural and functional features of the heart of 1-year-old *Rragc^S74N/+^* mice. Compared to wt mice of similar age, the decrease in heart weight was more pronounced in 1-year-old mice, as compared to young *Rragc^S74N/+^* mice, as measured by heart weight (Figure 2a) and by histological measurements of heart dimensions (Figure 2b and Figure S2a and S2b). The lack of a significant alteration in cardiomyocyte size (Figure S2c and S2d) points, as in the analysis of young individuals in Figure 1, to decreased cellularity. In contrast to results observed in the analysis of hearts of young mice, aged *Rragc^S74N/+^* mice showed fibrosis levels comparable to wild-type littermates (Figure S2e), indicating that fibrotic scars observed in hearts of young mice do not progress. Blood pressure was significantly increased in aged *Rragc*-mutant individuals (Figure 2c,^26^). Strikingly, echocardiographic analysis of old *Rragc^S74N/+^* mice (that had not succumbed during the perinatal period or later in life^26^) confirmed structural features consistent with a non-lethal DCM: decreased heart mass (Figure 2d), decreased IVS,d (Figure 2e), increased LVID,d/LV mass ratio (Figure 2f), and decreased aortic diameter (Figure S2f). In contrast to young mice, old *Rragc^S74N/+^* hearts present neither diastolic dysfunction neither exhibit aortic insufficiency (Figure 2g and 2h) and had no reduction in ejection fraction (Figure S2g). The analysis of surviving old *Rragc*-mutant mice suggest that survivors develop a milder cardiac condition with structural features of DCM but without compromised cardiac function.

**Figure 2.**
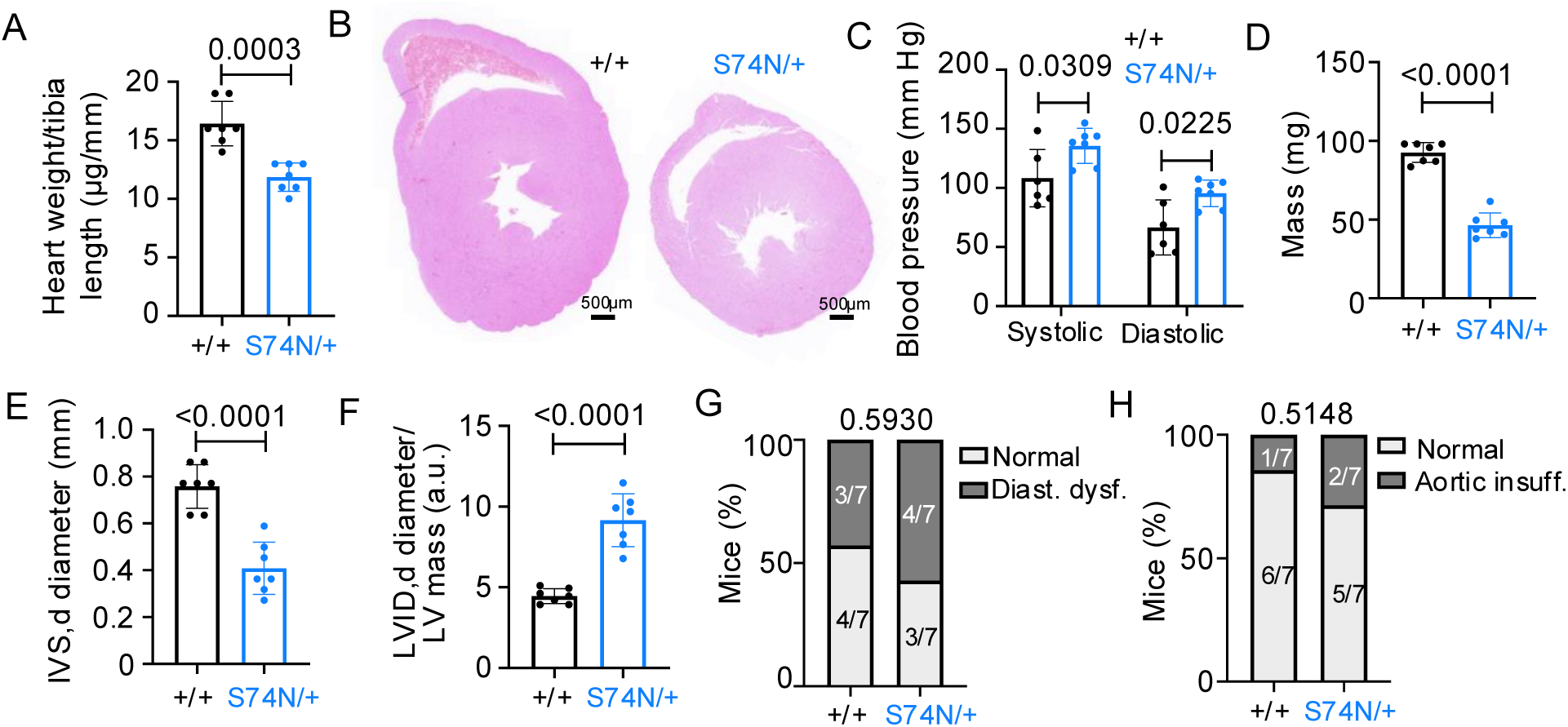
Old *RragcS74N/+* mice show DCM. **A.** Heart weight/tibia length ratio of 1.2 to 2.5 year-old *Rragc+/+* (n=7) and *RragcS74N/+* (n=7) mice. **B.** Representative images of heart size of 65 to 76 week-old *Rragc+/+* and *RragcS74N/+* mice. Scale bar 500μm. **C.** Blood pressure in 65-76 week-old *Rragc+/+* (n=6) and *RragcS74N/+* (n=7) mice. **D.** LV mass measured by echocardiographic analysis from 65-76 week-old in *Rragc+/+* (n=7) and *RragcS74N/+* (n=7) mice. **E.** IVS,d measured by echocardiographic analysis from 65-76 week-old in *Rragc+/+* (n=7) and *RragcS74N/+* (n=7) mice. **F.** LVID,d/LV mass measured by echocardiographic analysis from 65-76 week-old in *Rragc+/+* (n=7) and *RragcS74N/+* (n=7) mice. **G.** Diastolic dysfunction found by echocardiographic analysis from 65-76 week-old in *Rragc+/+* (n=7) and *RragcS74N/+* (n=7) mice. **H.** Aortic insufficiency found by echocardiographic analysis from 65-76 week-old in *Rragc+/+* (n=7) and *RragcS74N/+* (n=7) mice. In all panels data are presented as mean values ± D. Statistical significance was calculated by using unpaired two-tailed *t*-test in A, D, E, and F, 2-way ANOVA with Sidak’s multiple comparisons test in C, and chi-squared in G and H. p values are indicated in each panel.

### Transcriptome-wide interrogation of changes in *Rragc^S74N/+^* hearts reveals mitochondrial <u>dysfunction</u>

To identify molecular alterations caused by activating mutations in *Rragc* and to explain the early development of cardiac disease, we performed bulk RNAseq from neonatal wt and *Rragc^S74N/+^* hearts. This early time point was selected to assess changes in mRNA expression following the early-onset of DCM in the Long-Olsen-Distelmaier syndrome, given that hearts of *Rragc^S74N/+^*mice show alterations already at birth. Principal component analysis (PCA) clustered samples based on genotype (Figure 3a), suggesting robust changes in *Rragc^S74N/+^* vs. wt hearts. Gene set enrichment analysis (GSEA) revealed both expected and unanticipated signatures in *Rragc^S74N/+^*hearts. Consistently with the deposition of collagen fibers observed in *Rragc^S74N/+^*hearts (Figure 1e), several signatures related to collagen formation were largely enriched in *Rragc^S74N/+^* samples (Figure 3b). GSEA of *Rragc^S74N/+^* hearts showed signatures related to suppressed transcriptional activity of lysosomal biogenesis gene TFEB^25,32^ (Figure 3c). In addition, and importantly, we also observed a striking depletion of several metabolic signatures related to mitochondrial function (Figure 3b), and in particular, oxidative phosphorylation, mitochondrial fatty-acid oxidation (Figure 3d-e), mitochondria autophagy (Figure 3f) and respiratory chain complex (Figure 3b). This data prompt us to hypothesize that abnormal mitochondrial function and probably decreased mitochondrial respiration underlie the cardiac defects of *Rragc*-mutant mice.

**Figure 3.**
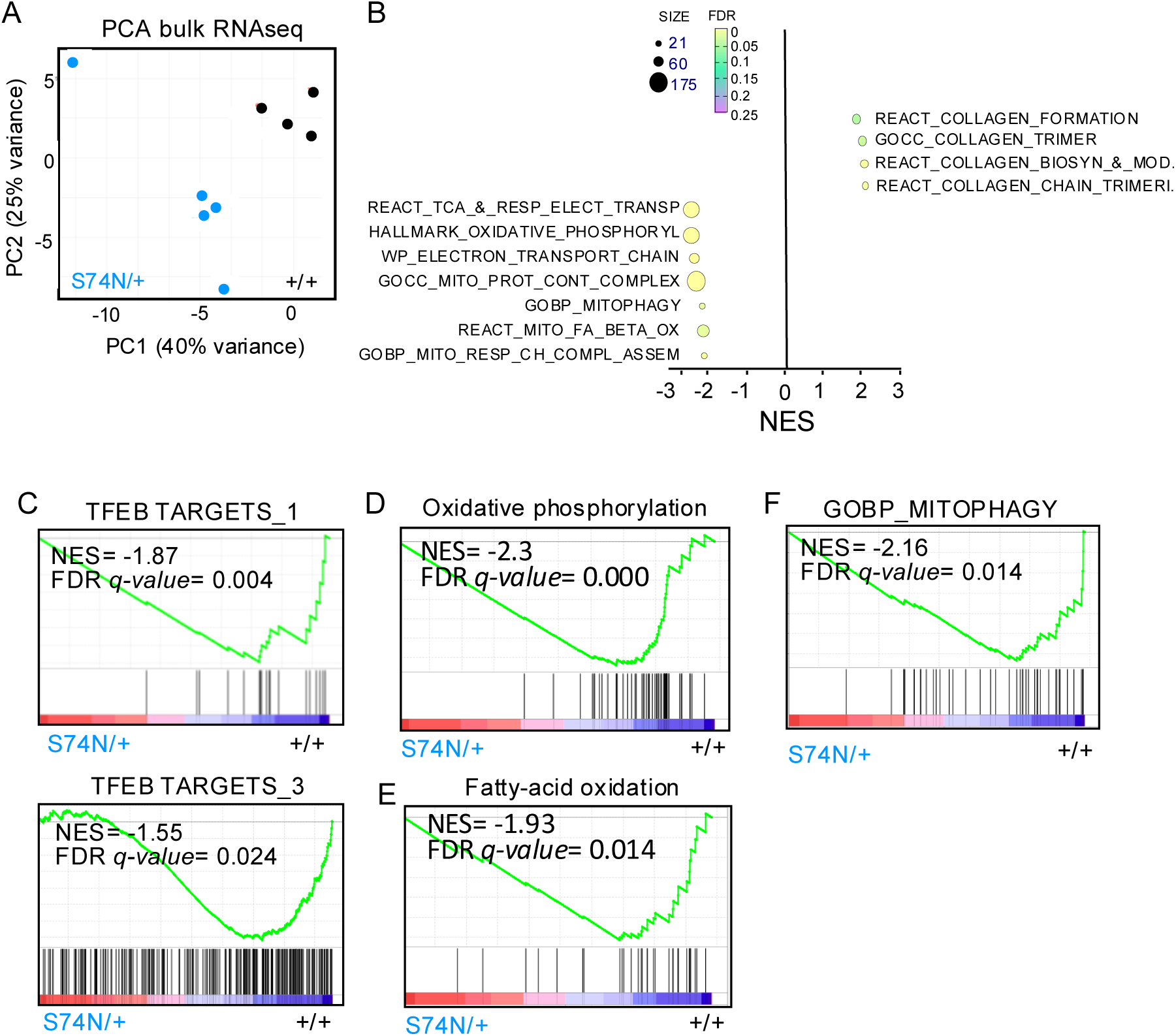
Transcriptomic analysis shows altered mitochondrial function in RagCS74N heart. **A.** Principal component analysis (PCA) from RNAseq performed from heart from neonatal *Rragc+/+* (n=4) and *RragcS74N/+* (n=5) mice. **B.** Bubble plot of the normalized enrichment score (NES) from the top gene sets enriched in hearts from neonatal *Rragc+/+* (n=4) versus *RragcS74N/+* (n=5) mice. **C.** Gene set enrichment analysis (GSEA) related to TFEB target genes in neonatal hearts from *Rragc+/+* (n=4) and *RragcS74N/+* (n=5). **D**. Gene set enrichment analysis (GSEA) of oxidative phosphorylation in neonatal hearts from *Rragc+/+* (n=4) and *RragcS74N/+* (n=5) **E**. Gene set enrichment analysis (GSEA) of fatty-acid oxidation in neonatal hearts from *Rragc+/+* (n=4) and *RragcS74N/+* (n=5) **F.** Gene set enrichment analysis (GSEA) of selective autophagy of mitochondria in neonatal hearts from *Rragc+/+* (n=4) and *RragcS74N/+* (n=5).

### Functional analysis of mitochondria from *Rragc^S74N/+^* cardiomyocytes shows abnormal <u>complex assembly</u>

To investigate the mitochondrial functional impairment caused by activating mutations in *Rragc*, we first analyzed the organization of the electron transport chain in complexes and supercomplexes by Blue Native page polyacrylamide gel electrophoresis of isolated cardiac mitochondria from adult *Rragc^+/+^* and *Rragc^S74N/+^*mice.

The results shows that CIV assembly into dimers was decreased in *Rragc^S74N/+^*hearts (Figure 4a-c), while there is no decrease of the amount of total Complex I, III and IV, strongly suggesting an abnormal molecular super-assembly of Complex IV (Figure S3a-d). Consistently with this idea, the mRNA levels of *Cox7a1*, encoding the subunit of Complex IV responsible for its dimerization^33^, were decreased in the bulk RNAseq analysis (STable 1 and Figure 4d) and validated by qPCR (Figure 4e).

**Figure 4.**
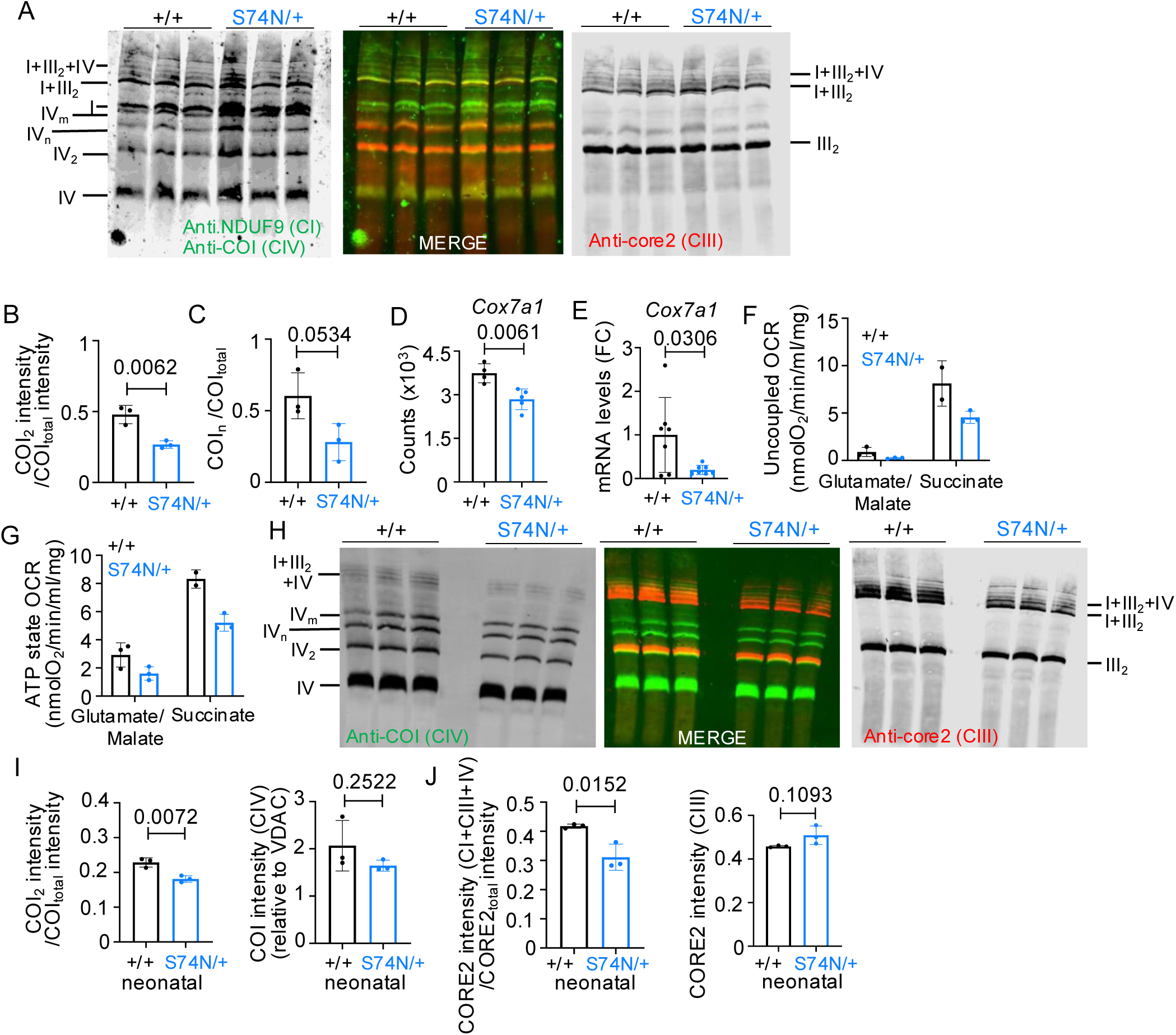
Defective respiratory complex assembly in hearts from adult and neonatal *RragcS74N/+* mice. **A.** Immunodetection of the indicated proteins (anti-COI for Complex IV and anti-NDUF9 for Complex I, anti-core2 for Complex III) after Blue Native PAGE of digitonin-solubilized mitochondria in adult heart from *Rragc+/+* (n=3) and *RragcS74N/+* (n=3). **B**. Quantification of the CIV assembled into supercomplexes from A. **C.** Quantification of complex IV multimers from A. **D.** Normalized counts for *Cox7a1* gene in RNAseq from hearts from *Rragc+/+* (n=4) and *RragcS74N/+* (n=5) mice shown in Figure 3. **E.** RT-qPCR of RNA from hearts obtained from 15-week-old *Rragc+/+* (n=7) and *RragcS74N/+* (n=7) male and female mice. Expression levels of is relative to the average level in wild-type mice. B2MG was used as housekeeping gene. **F.** Uncoupled oxygen consumption rate measured by oxygraphy in isolated mitochondria from hearts of 15-week-old *Rragc+/+* and *RragcS74N/+* mice. Mitochondria in each sample were obtained from a pool of three hearts. Glutamate/malate–driven respiration was measured in three technical replicates for both genotypes, whereas succinate-driven respiration was measured in two technical replicates for *Rragc+/+* and three technical replicates for *RragcS74N/+*. **G.** ATP-linked oxygen consumption rate measured under the same experimental conditions and using the same pooled mitochondrial preparations and technical replicates as in panel F. **H.** Immunodetection of the indicated proteins (anti-COI for Complex IV and anti-core2 for Complex III) after Blue Native PAGE of digitonin-solubilized mitochondria from neonatal heart from *Rragc+/+* (n=3, each pool of 5-6 hearts) and *RragcS74N/+* (n=3, each pool of 5-6 hearts). **I and J**. Quantification of CIV (I) and CIII_2_ (J) from figure H assembled into supercomplexes. In all panels data are presented as mean values ± SD. Statistical significance was calculated by using unpaired two-tailed *t*-test in B, C, D, E, I, and J. p values are indicated in each panel.

Neither enzymatic activity of complex IV (Figure S3e) nor respiratory control ratio in isolated mitochondria (Figure S3f) were affected in *Rragc^mut^* hearts. In contrast, and compatible with defective mitochondrial function, analysis oxygen consumption rate (OCR) from succinate-dependent respiration was lower in mitochondria pooled from *Rragc^S74N/+^* hearts (Figure 4f), and consistently, the maximum rate of mitochondrial oxygen consumption coupled to ATP production was decreased from both complex I and complex II-dependent respiration in isolated *Rragc^S74N/+^* mitochondria (Figure 4g). Mitochondrial DNA was significantly increased in mutant hearts (Fig S3g), pointing to a compensatory increase in mitochondrial mass typically observed in cases of mitochondrial dysfunction^34,35^. This increase could also reflect reduced mitochondrial turnover, consistently with the impaired mitophagy signature observed in RNAseq, and thus resulting in the accumulation of dysfunctional mitochondria and an overall rise in mtDNA content despite compromised respiratory capacity.

Finally, as the early phenotypic and transcriptomic changes are already present at birth, consistently with the early-onset nature of the Long-Olsen-Distelmaier syndrome, we assessed whether defects in mitochondrial electron transport organization were detectable in purified mitochondria from neonatal *Rragc*-mutant hearts. Indeed, purified mitochondria from neonatal *Rragc^S74N/+^* hearts exhibit decreased assembly of complex IV into dimers and decreased CIII assembled into CI+CIII_2_+CIV supercomplex (Figure 4h-j).

Altogether, the analysis of mitochondria from *Rragc*-mutant mice points to early abnormal mitochondrial complex assembly in hearts as a potential underlying mechanism of the early onset DCM in the Long-Olsen-Distelmaier syndrome.

## Discussion

We have modeled in mice key features of the Long-Olsen-Distelmaier syndrome by endogenous heterozygous expression of activating mutations in serine 74, orthologous to serine 75 in human *RRAGC* mutated in the disease. Although the mouse model did not reproduce cardiomegaly, reduced ejection fraction, or congenital septal defects such as atrial (ASD) and ventricular (VSD) septal defects, it displays a comparable DCM phenotype, characterized by decreased left ventricular mass and interventricular septal (IVS) diameter, along with increased LVID, cardiac insufficiency, and associated metabolic alterations.

This is not only the first report of genetic modeling of this DCM syndrome, but to our knowledge, a pioneer genetic system of a syndromic *Ragopathy*, recently coined as the group of human diseases caused by genetic deregulation of the Rag GTPases^22^. As such, this model provides a valuable experimental framework to dissect developmental consequences of Rag GTPase hyperactivation *in vivo*, which have remained largely unexplored.

Analysis of these mice revealed abnormal mitochondrial function, suggesting that early metabolic interventions should be considered, more so because of the early metabolic reprogramming that cardiomyocytes undergo after birth^36^ and the early onset of Long-Olsen-Distelmaier syndrome. Given the high energetic demand of cardiomyocytes, even subtle mitochondrial dysfunction is likely to have disproportionate effects on cardiac structure and performance, particularly during postnatal maturation. The mTORC1 inhibitor rapamycin stands out as an obvious and feasible intervention^23,37^, as it is already implemented in other syndromic diseases causing deregulated mTORC1 activity, such as tuberous sclerosis and Birt-Hogg-Dubé syndromes^38,39^. However, additional, metabolic interventions early in life may be also beneficial for Long-Olsen-Distelmaier patients. These could include strategies aimed at improving mitochondrial biogenesis, oxidative metabolism, or redox homeostasis, which warrant future investigation. Although longitudinal analyses starting from the fetal stages could provide additional insight into disease progression, the neonatal, adult and aging data presented here capture key early and established stages of the phenotype and provide meaningful mechanistic insight and potential intervention routes. Importantly, the consistency of phenotypes across these stages supports the notion that early molecular defects have lasting structural and functional consequences. Despite the large interspecies overlap, one difference between the human syndrome and the mouse model is that DCM is not lethal in the latter. Several explanations are possible, including the fact that the mutation is engineered in the germline and not somatic in mice. But as a matter of fact, *Rragc*-mutant mice are weaned at sub-mendelian ratios for reasons yet to be identified. Two complementary scenarios emerge: on the one hand, cardiac defects may underlie the partially-penetrant early lethality of *Rragc*-mutant mice, as cardiac abnormalities are already evident in *Rragc*-mutant neonates (Figure 1); on the other hand, it is tempting to speculate that underdiagnosed milder variants and manifestations of Long-Olsen-Distelmaier may exist in humans.

Based on the features that we have found we propose that Long-Olsen-Distelmaier syndrome may start to manifest during *in utero* development. This hypothesis is consistent with the early detection of cardiac and metabolic abnormalities and aligns with the known role of mTORC1 signaling in developmental growth control. Developmental cardiac damage associated to defects in mitochondria may result in fibrosis that compromises the relaxation capacity of the left ventricle, leading to altered ventricular pressure, ultimately resulting of aortic insufficiency and diastolic dysfunction. Such a pathogenic cascade provides a plausible mechanistic link between early metabolic impairment and progressive structural remodeling. Both pathologies share structural alterations in the heart, particularly to a decrease in the left ventricular mass and thickness, contributing to early-onset DCM, lethal in patients and milder and tolerated in mice. Together, these findings support a model in which mitochondrial and mTORC1-driven defects during cardiac development set the stage for long-term cardiomyopathy severity in a species-dependent manner.

## Methods

### Animal procedures

*Rragc^mut^* knock-in mouse models were generated in^24,26^. All animal procedures carried out at the CNIO were performed according to protocols approved by the CNIO-ISCIII Ethics Committee for Research and Animal Welfare (CEIyBA) and the Autonomous Community of Madrid (certificates PROEX285/15, PROEX015/18 and PROEX225.7/22). Mice were housed under specific pathogen free conditions at 22°C and with 12h dark/light cycles and fed with a standard chow diet (Harlan Teklad 2018). For 16h fasting experiments, mice were placed in a clean cage without access to food from 4pm to 8am next day. In each cage a maximum of two animals were placed. Under refeeding condition, mice had free access to food for 30min prior to sacrifice.

Pregnant females were injected with 1mg of progesterone (Sigma P0130) at embryonic days E17.5 and E18.5 to prevent premature delivery. At E19.5 females were euthanized and neonates were immediately obtained by C-section and placed on ice for sacrifice.

### Immunoblotting

For tissue extraction, a small piece of previously snap-frozen tissue was homogenized using FastPrep-24^TM^ 5 G instrument in ice-cold lysis buffer containing 50 mM HEPES (pH 7.4), 40 mM NaCl, 2 mM EDTA, 1.5 mM sodium orthovanadate, 50 mM NaF, 10 mM pyrophosphate, 10 mM glycerophosphate, 1% Triton X-100 and one tablet of EDTA-free complete protease inhibitors (Roche #11836153001) per 25 ml. Tissue lysates were cleared by centrifugation at maximum speed for 10 min at 4°C. Protein content was measured with BCA Protein Assay (Thermo Scientific™ #23222). Protein extracts were denatured by adding sample buffer and boiling for 5 min, resolved by SDS-PAGE and analyzed by immunoblotting. Results were visualized using Odyssey Infrared Imaging System (Application software version 3.0.30) LI-COR Biosciences. The following primary antibodies were used: P-S235/236-S6 (Cell Signaling Technology #2211), S6 (Cell Signaling Technology #2217), RagC (Cell Signaling Technology #9480) and anti-vinculin (Sigma #V9131).

### Histology and Immunohistochemical staining

Tissue samples intended for hematoxylin and eosin (H&E) and Sirius Red staining were fixed in 10% neutral buffered formalin (4% formaldehyde in solution), embedded in paraffin, and sectioned at a thickness of 3 μm. Sections were mounted on Superfrost Plus™ slides and dried overnight. Prior to staining, slides were deparaffinized in xylene and rehydrated through a graded ethanol series. Subsequently, sections were stained with either H&E for general histological evaluation or Sirius Red to assess collagen deposition and fibrosis. For immunofluorescence staining against Wheat Germ Agglutinin, sections were fixed 10 min with 4% PFA (Electron Microscopy Sciences #15710) in PBS at room temperature and then wash three times 5 min with PBS. Sections were blocked/permeabilized 1h at room temperature in PBS with 10% Donkey Serum (Jackson ImmunoResearch #017-000-121), 10% FBS (Hyclone #SV30160.03) and 0.5% Triton (Sigma #T9284). Primary antibody (Wheat Germ Agglutinin, Alexa Fluor 488 conjugate (Thermo Fisher #W11261) dilution: 1/500) was diluted in blocking/permeabilization buffer and incubated overnight at 4°C. This step was followed by three washes in PBS of 5 min each and incubation for 2h with conjugated secondary antibodies (1:300) and DAPI (Sigma #D9542) at room temperature. After three washes of 5 min in PBS, cells were mounted with Prolong Gold antifade reagent (Cell Signaling Technology #9071).

### Quantification of fibrosis

Fibrosis in heart tissue was quantified from Sirius Red-stained sections using ImageJ. The fibrotic area was measured and normalized to the total tissue area.

### Echocardiography

Biomedical Imaging has been conducted at the Advanced Imaging Unit of CNIC. Ultrahigh resolution echocardiography was performed by operator blind to the origin of the animals with a linear 30-MHz transducer (Vevo F2 VisualSonics Inc). Mice were lightly anesthetized with 1 to 1.5% isoflurane in 100% oxygen and placed on a heated table. Heart rate (HR) was monitored and anesthesia delivery was adjusted to maintain an HR of 450 ± 50 beats/min. Two-dimensional and M-mode (MM) echocardiography was performed in parasternal long– and short axis views. Left ventricle diastolic function was assessed by spectral Doppler echocardiography analyzing early (E wave) and late (A wave) velocities of mitral flow, also pulse-wave Doppler of descending aorta was recorded in the longitudinal view. Finaly, to obtain Pulse Wave Velocity (PWV), a two-dimensional image of the ascending aorta, aortic arch and descending aorta was acquired. Pulsed Doppler was then acquired in the ascending and descending aorta.

### Blood pressure measurement

Blood pressure was measured using a noninvasive tail-cuff method with the CODA system (CODA-HT2, Kent Scientific Corporation)^40^. Mice were trained for three consecutive days before taking the final measurements.

### Bulk RNA sequencing

#### RNA extraction

For total RNA extraction, a previously snap-frozen whole neonatal heart was homogenized in Trizol (Invitrogen #15596026) using the suggested homogenization protocols of the FastPrep machine (FastPrep-24^TM^ 5 G). For RNA extraction, RNeasy Mini Kit (Qiagen #74106) was used, following manufacturer’s instructions. Total RNA samples (500ng) were converted into cDNA sequencing libraries with the QuantSeq 3‘ mRNA-Seq V2 Library Prep Kit (FWD) for Illumina (Lexogen #191). Briefly, library generation is initiated by reverse transcription with oligodT priming, followed by a random-primed second strand synthesis. Primers from both steps contain Illumina-compatible sequences. Libraries were completed by PCR and sequenced on an Illumina NextSeq 550 instrument (with v2.5 reagent kits) by following manufacturer’s protocols. This library preparation kit generates directional libraries stranded in the sense orientation, the read1 (the only read in single read format) has the sense orientation. Raw images generated by the sequencer are submitted to analysis, per-cycle basecalling and quality score assignment with Illumina’s Real Time Analysis integrated primary analysis software (RTA v2). Conversion of BCL (base calls) binary files to FASTQ format is subsequently performed with bcl2fastq2 (Illumina). FASTQ files obtained from RNA sequencing were subjected to quality control using FastQC (version 0.11.9)^41^. PolyA tails and adapter sequences were removed using cutadapt (version 4.0)^41^. The resulting reads were then processed using the cluster_RNAseq pipeline (https://github.com/cnio-bu/cluster_rnaseq. In this process), the reads were aligned to the GRCm39 version of the mouse reference genome using STAR (version 2.7.11a)^42^. Genomic feature counts were obtained using featureCounts^43^. Differential expression analysis was performed using DESeq2 (version 1.30.1)^44^. Differentially expressed genes were identified based on an adjusted p-value threshold of 0.05 or lower. Gene set enrichment analysis was performed using GSEA (version 4.3.2)^45^. The matrix used for the analysis contains normalized values derived from the results of the differential expression analysis. To generate the matrix, Ensembl IDs were converted to gene symbols using biomaRt (version 2.54.0) from the R bioconductor package^46^, utilizing the mmusculus_gene_ensembl dataset and ensembl database version 110. For each gene symbol, the maximum absolute value was selected. Non-coding RNAs were excluded from further analysis. The parameters used for the GSEA analysis were the default ones except for the following: –collapse No_Collapse, –permute gene_set, –plot_top_x 100, and –set_min 10. Gene sets from MSigDB v2023.2.Mm were utilized for the analysis. Specifically, the following gene sets from the mouse collection in MSigDB were used: curated gene sets of canonical pathways, gene ontology gene sets, cell type signature gene sets and hallmark gene sets.

### Quantitative PCR

Total RNA samples (500 ng) were retro-transcribed using the High Capacity cDNA Reverse Transcription Kit (Applied Biosystems #4368814) following manufacturer’s instructions. For Cox7a1 assessment primers used are F: AAACCGTGTGGCAGAGAAGCAG R: CCCAAGCAGTATAAGCAGTAGGC. Quantitative real-time PCR was run in triplicates using GoTaq qPCR Master Mix (Promega #A6001) with a reaction volume of 10 μL in a QuantStudio 6 Flex Real-Time PCR System thermocycler (Applied Biosystems). Data were analyzed by the change-in-threshold (2^−ΔΔCT^) method, using β-actin as “housekeeping” reference gene for mouse samples. Results are represented as fold changes relative to the mean expression levels of the control condition.

### Isolation of mitochondria

Mitochondria from the heart were isolated according to the previously described method^47^. Heart tissue was disaggregated with blade and homogenized in solution containing sucrose 0,32M, EDTA 1 mM, TRIS-HCL 10mM, pH 7,4, using a Potter-Elvehjem at 600 rpm. After an initial centrifugation at 600*g* at 4°C, 10 min, the supernatant was collected and centrifuged at 10.000*g* 4°C 10 min, twice. The pellet obtained was resuspended in Sucrose 0,32M, EDTA 1 mM, TRIS-HCL 10mM, pH 7,4, and mitochondrial protein concentration was quantified by Bradford assay (Bio-Rad #5000006).

### Mitochondrial membrane solubilization

To detect respiratory chain complexes and supercomplexes, mitochondria were suspended at 10 mg/ml in 1,5M aminocaproic acid, 50 mM Bis-Tris/HCl pH 8, followed by solubilization with digitonin on ice at a final concentration of 4% for 5 min or with DDM (dodecyl-maltoside) at final concentration of 1%. After centrifugation at 16,000*g* for 20 min at 4°C, the supernatant was collected and prepared with Serva Blue G dye (Serva Scientist # 35050.03), stock 5%.

### Blue native gel electrophoresis

Samples were separated in non-denaturing conditions on a 3–12% gradient gel using a peristaltic pump and casting the gel into Biorad Mini protein system^48^. Gels were electroblotted onto Hybond-P-polyvinylidene fluoride (PVDF) membranes (GE Healthcare) and immunoblotted with specific antibodies against the different subunits of the respiratory complexes: anti-UQCRC2 or CORE2 (ProteinTech #14742-1-AP), anti-MTCO1 (Invitrogen #459600), anti-NDUFA9 (Abcam #ab14713) The secondary antibodies were conjugated to AlexaFluor 680 (Invitrogen #A21076) or AF800 (Invitrogen #SA5-10176) and acquired with the ODYSSEY Infrared Imaging System (LI-COR).

### Mitochondrial DNA content

Mitochondrial DNA content was measured as described previously^49^, with some with modifications. DNA extraction from heart was performed following instructions in DNeasy kit (QIAGEN #69504), and using 100μl of elution buffer in the last step. Nuclear and mitochondrial DNA were amplified by quantitative PCR with 10ng of total DNA as template, using specific primers for mitochondrial gene mtCOI (mtCO1 Forward: CTGAGCGGGAATAGTGGGTA; mtCO1 Reverse: TGGGGCTCCGATTATTAGTG) and nuclear gene SDHA (nSDHA Forward: TACTACAGCCCCAAGTCT; nSDHA Reverse: T TGGACCCATCTTCTATGC) and GoTaq qPCR Master Mix (Promega #A6001). Mitochondrial content was normalized to nuclear DNA and was calculated using the equation: ΔC_T_ = (nucDNA C_T_ – mtDNA C_T_). Relative mitochondrial DNA copies = 2 × 2^ΔCT^.

### Respiration assays

Respiration assays were performed using the Oxytherm-R (HansTech) equipment, that allows measurement of the respiration capacity using different substrates. For the experiments, 3 hearts from mice of each genotype were pooled together and mitochondria were isolated as described above. The respiration assays were performed with 300 μg mitochondria using as substrates: glutamate-malate at final concentration 10mM (allows the entrance of electrons from the ETC-Complex I), succinate, at final concentration 10mM (allows the entrance of electrons from the ETC-Complex II), or malate + palmitoyl-carnitine, at final concentration 0.5mM and 0.05mM respectively (coupling oxygen consumption with β-oxidation)^50^.

### Cytochrome c oxidase activity

Cytochrome c oxidase (Complex IV) activity was measured spectrophotometrically by monitoring the oxidation of reduced cytochrome c. Reduced cytochrome c was prepared using sodium dithionite, and excess reductant was removed prior to use. The assay was performed at [25 °C or 37 °C] in 50 mM potassium phosphate buffer (pH 7.0) containing 0.05% n-dodecyl-β-D-maltoside and 10–50 µM reduced cytochrome c.

The reaction was initiated by the addition of isolated mitochondria and the decrease in absorbance at 550 nm was recorded for 1–2 minutes using a UV–visible spectrophotometer. Enzymatic activity was calculated from the linear portion of the trace using an extinction coefficient of ε₅₅₀ = 19.1 mM⁻¹·cm⁻¹ and expressed as nmol cytochrome c oxidized·min⁻¹·mg⁻¹ protein. Background oxidation was subtracted, and assay specificity was confirmed by inhibition with 1 mM potassium cyanide.

### Statistical analysis

The Ν, indicating the total number of animals per group is indicated in each figure legend. Statistical analysis was carried out with Prism 8 (GraphPad). Unpaired t-test was used for single comparisons. One-way ANOVA with Tukey’s multiple comparisons test was used for continuous variables. All experiments that include a second variable were analyzed using Two-way ANOVA with Sidak’s multiple comparison post-test. Chi-squared test was used when comparing contingency data. Εrror bars in all panels depict standard deviation (SD).

### Data availability

Sequence data that support the findings of this study have been deposited in GEO, with the accession code GSE302581 in private mode (during the review process). To review GEO accession GSE302581: https://urldefense.com/v3/ https://www.ncbi.nlm.nih.gov/geo/query/acc.cgi?acc=GSE302581 <u>;!!D9dNQwwGXtA!XNZJnSiEg4Hhg_DYbzYBz-6395YiclSMlhVW2qa8HbXuZ8CynoexiSASgl1hpxn86ugLOHKjwfZ7kizKKA$</u>

Enter token **epqtseeipnontsp** into the box. Additional data are available from the corresponding authors upon request.

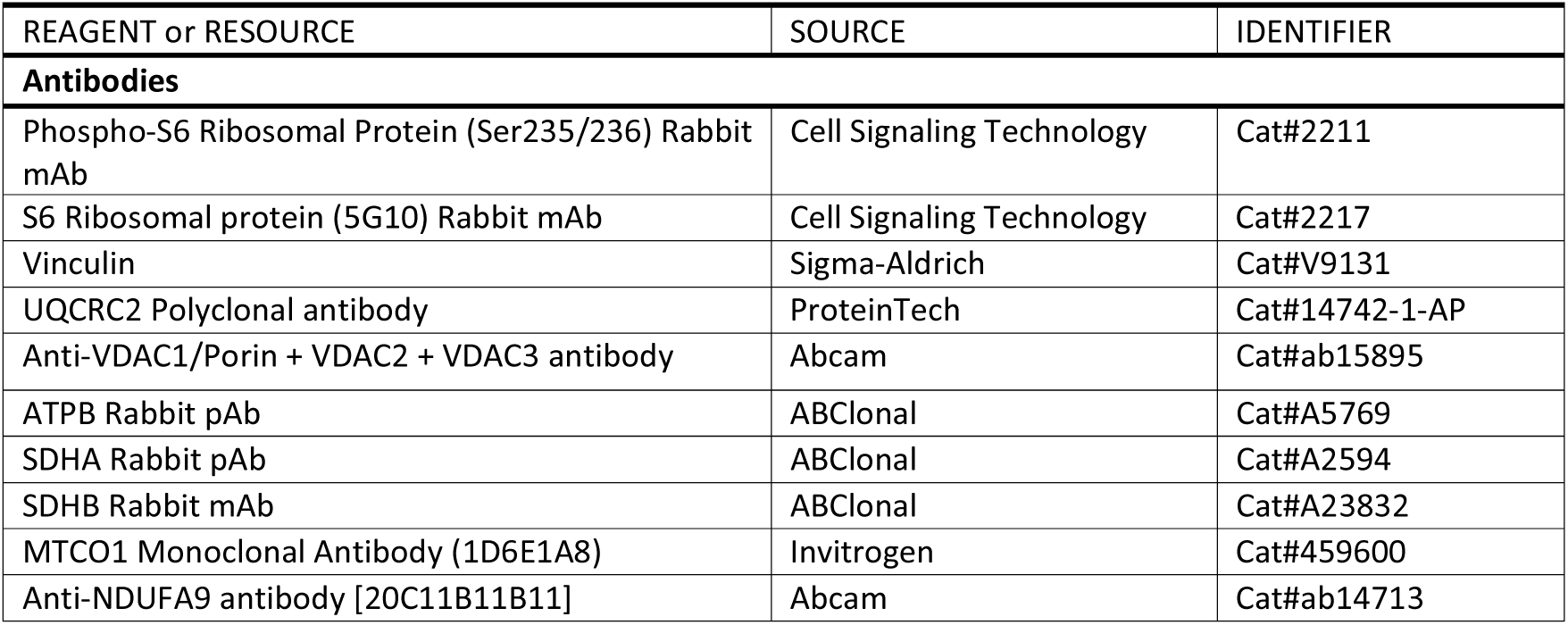
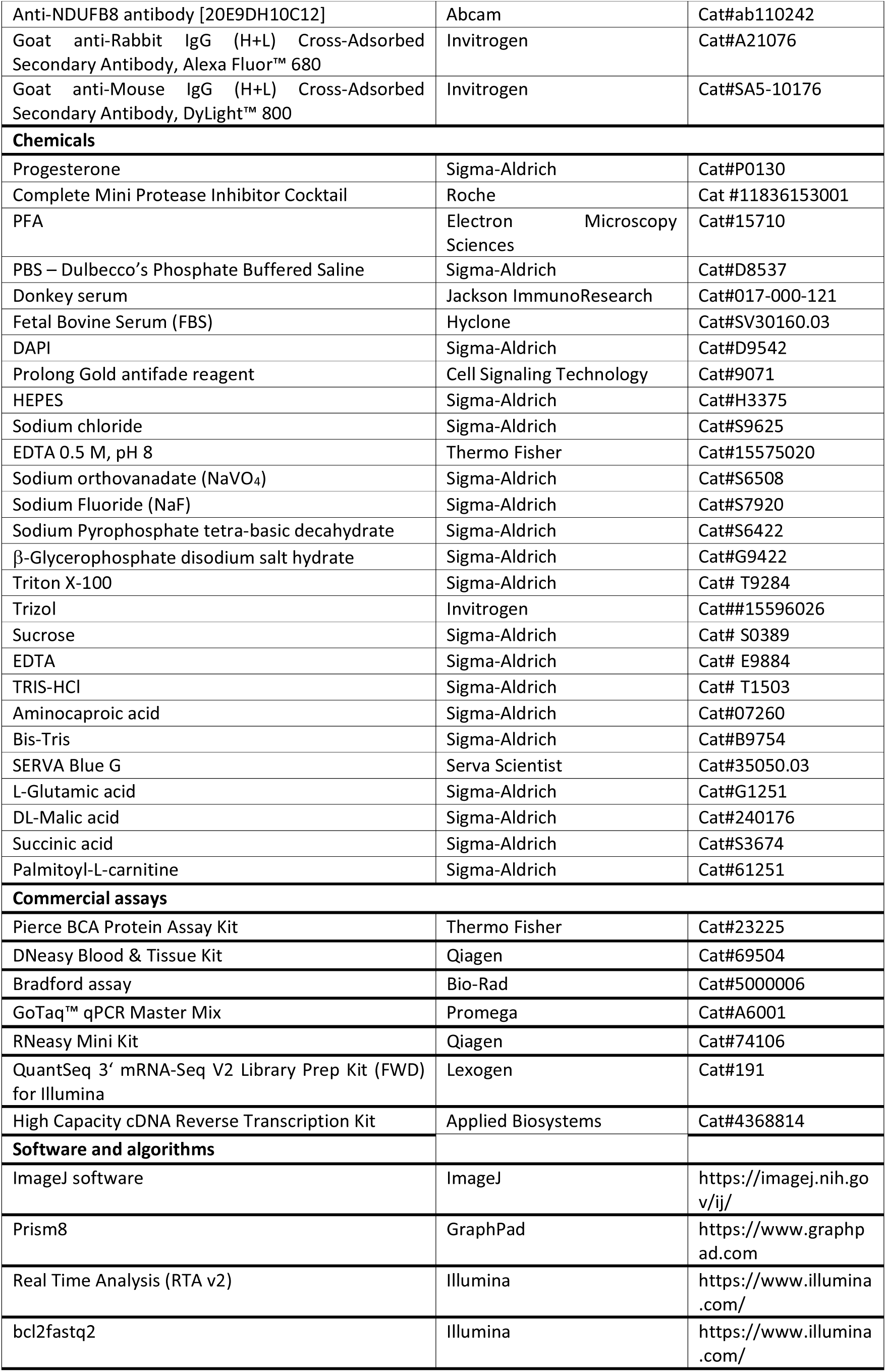
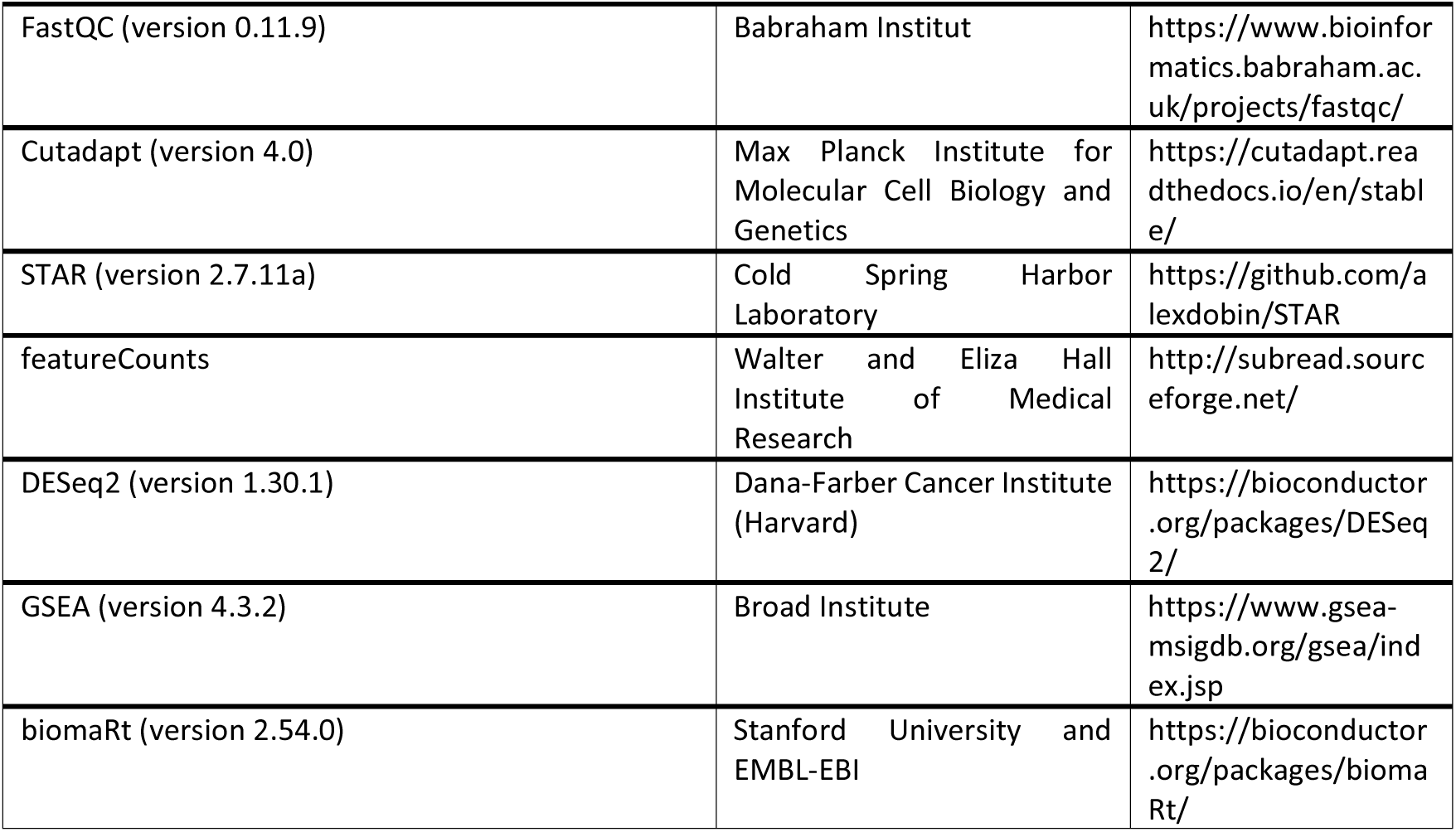
Key Resources Table.

## Acknowledgements

We thank the CNIO Histopathology, Animal Facility and Genomics Core Units for excellent technical support. Biomedical imaging has been conducted at the Advanced Imaging Unit of the CNIC (Centro Nacional de Investigaciones Cardiovasculares Carlos III), Madrid, Spain. Research was supported by the RETOS projects Programme of Spanish Ministry of Science, Innovation and Universities, Spanish State Research Agency, co-funded by the European Regional Development Fund (grant PID2019-104012RB-I00 AND and PID2022-136413OB-I00), EU-H2020 Programme (ERC-2014-STG-638891), FERO Grant for Research in Oncology and LaCaixa Banking Foundation (LCF/PR/HR21/0046). S.C. is supported by the CNS2023-143646 founded by MICIU/AEI/10.13039/501100011033 and European Union NextGenerationEU/PRTR and Grant PID2023-148516OB-I00 funded by MCIN/AEI/ 10.13039/501100011033. A.B.P.-G, L d.P.-R., E.S.G. and L.F. are recipients of Ayudas de contratos predoctorales para la formación de doctores from MICIU/AEI (BES-2017-081381, PRE-2019-090891, and PRE2020-095127, PRE2022-000052). A.M.S is supported by La Caixa Foundation (HR24-00581), Generalitat Valenciana PROMETEO CIPROM/2022/48 project and PMP21/00057 IMPACT-2021, funded by the Instituto de Salud Carlos III (ISCIII). A.E. is an EMBO Young Investigator

## Author Contributions

A.B.P.-G. and E.S.G. performed most experimental work and analyzed data with support from L.d.P.-R., A.S., A.O.-M., and L.F.. A.Sag. and S.C. performed mitochondrial analyses and contributed to data conceptualization. A.M.S., L.L.V, A.C.A., G.S. contributed electrocardiographic analyses and data analysis and conceptualization. E.C. performed pathology analysis. E.P.Y. performed bioinformatics analysis. A.E. conceived and supervised the study, analyzed the data, wrote the manuscript and secured funding. All authors read and commented on the manuscript and figures.

## References

1. Pinto, Y.M., Elliott, P.M., Arbustini, E., Adler, Y., Anastasakis, A., Böhm, M., Duboc, D., Gimeno, J., De Groote, P., Imazio, M., et al. (2016). Proposal for a revised definition of dilated cardiomyopathy, hypokinetic non-dilated cardiomyopathy, and its implications for clinical practice: A position statement of the ESC working group on myocardial and pericardial diseases. Eur Heart J 37. 10.1093/eurheartj/ehv727.

2. Seferović, P.M., Polovina, M., Bauersachs, J., Arad, M., Gal, T. Ben, Lund, L.H., Felix, S.B., Arbustini, E., Caforio, A.L.P., Farmakis, D., et al. (2019). Heart failure in cardiomyopathies: a position paper from the Heart Failure Association of the European Society of Cardiology. Eur J Heart Fail 21. 10.1002/ejhf.1461.

3. McNally, E.M., and Mestroni, L. (2017). Dilated cardiomyopathy: Genetic determinants and mechanisms. Preprint, https://doi.org/10.1161/CIRCRESAHA.116.309396 10.1161/CIRCRESAHA.116.309396.

4. Schultheiss, H.-P., Fairweather, D., Caforio, A.L.P., Escher, F., Hershberger, R.E., Lipshultz, S.E., Liu, P.P., Matsumori, A., Mazzanti, A., McMurray, J., et al. (2019). Dilated cardiomyopathy. Nat Rev Dis Primers 5, 32. 10.1038/s41572-019-0084-1.

5. Mozaffarian, D., Benjamin, E.J., Go, A.S., Arnett, D.K., Blaha, M.J., Cushman, M., Das, S.R., de Ferranti, S., Després, J.-P., Fullerton, H.J., et al. (2016). Heart Disease and Stroke Statistics—2016 Update. Circulation 133. 10.1161/cir.0000000000000350.

6. Eldemire, R., Mestroni, L., and Taylor, M.R.G. (2024). Genetics of Dilated Cardiomyopathy. Preprint, https://doi.org/10.1146/annurev-med-052422-020535 10.1146/annurev-med-052422-020535.

7. Zheng, S.L., Henry, A., Cannie, D., Lee, M., Miller, D., McGurk, K.A., Bond, I., Xu, X., Issa, H., Francis, C., et al. (2024). Genome-wide association analysis provides insights into the molecular etiology of dilated cardiomyopathy. Nat Genet 56, 2646–2658. 10.1038/s41588-024-01952-y.

8. Jurgens, S.J., Rämö, J.T., Kramarenko, D.R., Wijdeveld, L.F.J.M., Haas, J., Chaffin, M.D., Garnier, S., Gaziano, L., Weng, L.-C., Lipov, A., et al. (2024). Genome-wide association study reveals mechanisms underlying dilated cardiomyopathy and myocardial resilience. Nat Genet 56, 2636–2645. 10.1038/s41588-024-01975-5.

9. Martinez, H.R., Beasley, G.S., Miller, N., Goldberg, J.F., and Jefferies, J.L. (2021). Clinical Insights Into Heritable Cardiomyopathies. Preprint, 10.3389/fgene.2021.663450 https://doi.org/10.3389/fgene.2021.663450.

10. Reijnders, M.R.F., Seibt, A., Brugger, M., Lamers, I.J.C., Ott, T., Klaas, O., Horváth, J., Rose, A.M.S., Craghill, I.M., Brunet, T., et al. (2023). De novo missense variants in RRAGC lead to a fatal mTORopathy of early childhood. Genetics in Medicine 25. 10.1016/j.gim.2023.100838.

11. Long, P.A., Zimmermann, M.T., Kim, M., Evans, J.M., Xu, X., and Olson, T.M. (2016). De novo RRAGC mutation activates mTORC1 signaling in syndromic fetal dilated cardiomyopathy. Hum Genet. 10.1007/s00439-016-1685-3.

12. Liu, G.Y., and Sabatini, D.M. (2020). mTOR at the nexus of nutrition, growth, ageing and disease. Preprint, 10.1038/s41580-019-0199-y https://doi.org/10.1038/s41580-019-0199-y.

13. Efeyan, A., Comb, W.C., and Sabatini, D.M. (2015). Nutrient-sensing mechanisms and pathways. Nature 517, 302–310.

14. Shimobayashi, M., and Hall, M.N. (2014). Making new contacts: the mTOR network in metabolism and signalling crosstalk. Nat Rev Mol Cell Biol 15, 155–162. 10.1038/nrm3757.

15. Saxton, R.A., and Sabatini, D.M. (2017). mTOR Signaling in Growth, Metabolism, and Disease. Preprint, 10.1016/j.cell.2017.02.004 https://doi.org/10.1016/j.cell.2017.02.004.

16. Lawrence, R.E., and Zoncu, R. (2019). The lysosome as a cellular centre for signalling, metabolism and quality control. Preprint, https://doi.org/10.1038/s41556-018-0244-7 10.1038/s41556-018-0244-7.

17. Sancak, Y., Peterson, T.R., Shaul, Y.D., Lindquist, R.A., Thoreen, C.C., Bar-Peled, L., and Sabatini, D.M. (2008). The Rag GTPases bind raptor and mediate amino acid signaling to mTORC1. Science (1979) 320, 1496–1501. https://doi.org/1157535 [pii] 10.1126/science.1157535.

18. Okosun, J., Wolfson, R.L., Wang, J., Araf, S., Wilkins, L., Castellano, B.M., Escudero-Ibarz, L., Al Seraihi, A.F., Richter, J., Bernhart, S.H., et al. (2015). Recurrent mTORC1-activating RRAGC mutations in follicular lymphoma. Nat Genet. 10.1038/ng.3473.

19. Ying, Z.X., Jin, M., Peterson, L.F., Bernard, D., Saiya-Cork, K., Yildiz, M., Wang, S., Kaminski, M.S., Chang, A.E., Klionsky, D.J., et al. (2016). Recurrent mutations in the MTOR regulator RRAGC in follicular lymphoma. In Clinical Cancer Research 10.1158/1078-0432.CCR-16-0609.

20. Green, M.R., Kihira, S., Liu, C.L., Nair, R. V, Salari, R., Gentles, A.J., Irish, J., Stehr, H., Vicente-Duenas, C., Romero-Camarero, I., et al. (2015). Mutations in early follicular lymphoma progenitors are associated with suppressed antigen presentation. Proc Natl Acad Sci U S A 112, E1116–25. 10.1073/pnas.1501199112.

21. Sambri, I., Ferniani, M., Campostrini, G., Testa, M., Meraviglia, V., de Araujo, M.E.G., Dokládal, L., Vilardo, C., Monfregola, J., Zampelli, N., et al. (2023). RagD auto-activating mutations impair MiT/TFE activity in kidney tubulopathy and cardiomyopathy syndrome. Nat Commun 14. 10.1038/s41467-023-38428-2.

22. Sambri, I., Ferniani, M., and Ballabio, A. (2024). Ragopathies and the rising influence of RagGTPases on human diseases. Nat Commun 15, 5812. 10.1038/s41467-024-50034-4.

23. Kim, M., Lu, L., Dvornikov, A. V., Ma, X., Ding, Y., Zhu, P., Olson, T.M., Lin, X., and Xu, X. (2021). Tfeb overexpression, not mtor inhibition, ameliorates ragcs75y cardiomyopathy. Int J Mol Sci 22. 10.3390/ijms22115494.

24. Ortega-Molina, A., Deleyto-Seldas, N., Carreras, J., Sanz, A., Lebrero-Fernández, C., Menéndez, C., Vandenberg, A., Fernández-Ruiz, B., Marín-Arraiza, L., de la Calle Arregui, C., et al. (2019). Oncogenic Rag GTPase signalling enhances B cell activation and drives follicular lymphoma sensitive to pharmacological inhibition of mTOR. Nat Metab. 10.1038/s42255-019-0098-8.

25. Napolitano, G., Di Malta, C., Esposito, A., de Araujo, M.E.G., Pece, S., Bertalot, G., Matarese, M., Benedetti, V., Zampelli, A., Stasyk, T., et al. (2020). A substrate-specific mTORC1 pathway underlies Birt–Hogg–Dubé syndrome. Nature. 10.1038/s41586-020-2444-0.

26. Ortega-Molina, A., Lebrero-Fernandez, C., et al, and Efeyan, A. (2024). A mild increase in nutrient signaling to mTORC1 in mice leads to parenchymal damage, myeloid inflammation and shortened lifespan. Nat Aging, 1102–1120. 10.1038/s43587-024-00635-x.

27. Efeyan, A., Zoncu, R., Chang, S., Gumper, I., Snitkin, H., Wolfson, R.L., Kirak, O., Sabatini, D.D., and Sabatini, D.M. (2013). Regulation of mTORC1 by the Rag GTPases is necessary for neonatal autophagy and survival. Nature 493, 679–683.

28. Polak, P., and Hall, M.N. (2009). mTOR and the control of whole body metabolism. Curr Opin Cell Biol 21, 209–218. 10.1016/j.ceb.2009.01.024.

29. Loewith, R., Jacinto, E., Wullschleger, S., Lorberg, A., Crespo, J.L., Bonenfant, D., Oppliger, W., Jenoe, P., and Hall, M.N. (2002). Two TOR complexes, only one of which is rapamycin sensitive, have distinct roles in cell growth control. Mol Cell 10, 457–468. S1097276502006366 [pii].

30. Ersching, J., Efeyan, A., Mesin, L., Jacobsen, J.T., Pasqual, G., Grabiner, B.C., Dominguez-Sola, D., Sabatini, D.M., and Victora, G.D. (2017). Germinal Center Selection and Affinity Maturation Require Dynamic Regulation of mTORC1 Kinase. Immunity. 10.1016/j.immuni.2017.06.005.

31. Kim, Y.C., Park, H.W., Sciarretta, S., Mo, J.S., Jewell, J.L., Russell, R.C., Wu, X., Sadoshima, J., and Guan, K.L. (2014). Rag GTPases are cardioprotective by regulating lysosomal function. Nat Commun. 10.1038/ncomms5241.

32. Wada, S., Neinast, M., Jang, C., Ibrahim, Y.H., Lee, G., Babu, A., Li, J., Hoshino, A., Rowe, G.C., Rhee, J., et al. (2016). The tumor suppressor FLCN mediates an alternate mTOR pathway to regulate browning of adipose tissue. Genes Dev. 10.1101/gad.287953.116.

33. García-Poyatos, C., Arora, P., Calvo, E., Marques, I.J., Kirschke, N., Galardi-Castilla, M., Lembke, C., Meer, M., Fernández-Montes, P., Ernst, A., et al. (2024). Cox7a1 controls skeletal muscle physiology and heart regeneration through complex IV dimerization. Dev Cell 59, 1824–1841.e10. 10.1016/j.devcel.2024.04.012.

34. Filograna, R., Mennuni, M., Alsina, D., and Larsson, N.G. (2021). Mitochondrial DNA copy number in human disease: the more the better? Preprint, https://doi.org/10.1002/1873-3468.14021 10.1002/1873-3468.14021.

35. Castellani, C.A., Longchamps, R.J., Sun, J., Guallar, E., and Arking, D.E. (2020). Thinking outside the nucleus: Mitochondrial DNA copy number in health and disease. Preprint, https://doi.org/10.1016/j.mito.2020.06.004 10.1016/j.mito.2020.06.004.

36. Girard, J., Ferre, P., Pegorier, J.P., and Duee, P.H. (1992). Adaptations of glucose and fatty acid metabolism during perinatal period and suckling-weaning transition. Physiol Rev 72, 507–562.

37. McMullen, J.R., Sherwood, M.C., Tarnavski, O., Zhang, L., Dorfman, A.L., Shioi, T., and Izumo, S. (2004). Inhibition of mTOR signaling with rapamycin regresses established cardiac hypertrophy induced by pressure overload. Circulation 109. 10.1161/01.CIR.0000130641.08705.45.

38. Geilswijk, M., Genuardi, M., Woodward, E.R., Nightingale, K., Huber, J., Madsen, M.G., Liekelema – van der Heij, D., Lisseman, I., Marlé-Ballangé, J., McCarthy, C., et al. (2024). ERN GENTURIS clinical practice guidelines for the diagnosis, surveillance and management of people with Birt-Hogg-Dubé syndrome. European Journal of Human Genetics 32, 1542–1550. 10.1038/s41431-024-01671-2.

39. Henske, E.P., Jóźwiak, S., Kingswood, J.C., Sampson, J.R., and Thiele, E.A. (2016). Tuberous sclerosis complex. Nat Rev Dis Primers 2, 16035. 10.1038/nrdp.2016.35.

40. Simon Andrews (2020). Babraham Bioinformatics – FastQC A Quality Control tool for High Throughput Sequence Data. Soil 5.

41. Martin, M. (2011). Cutadapt removes adapter sequences from high-throughput sequencing reads. EMBnet J 17. 10.14806/ej.17.1.200.

42. Dobin, A., Davis, C.A., Schlesinger, F., Drenkow, J., Zaleski, C., Jha, S., Batut, P., Chaisson, M., and Gingeras, T.R. (2013). STAR: Ultrafast universal RNA-seq aligner. Bioinformatics 29. 10.1093/bioinformatics/bts635.

43. Liao, Y., Smyth, G.K., and Shi, W. (2014). FeatureCounts: An efficient general purpose program for assigning sequence reads to genomic features. Bioinformatics 30. 10.1093/bioinformatics/btt656.

44. Love, M.I., Huber, W., and Anders, S. (2014). Moderated estimation of fold change and dispersion for RNA-seq data with DESeq2. Genome Biol. 10.1186/s13059-014-0550-8.

45. Subramanian, A., Tamayo, P., Mootha, V.K., Mukherjee, S., Ebert, B.L., Gillette, M.A., Paulovich, A., Pomeroy, S.L., Golub, T.R., Lander, E.S., et al. (2005). Gene set enrichment analysis: A knowledge-based approach for interpreting genome-wide expression profiles. Proceedings of the National Academy of Sciences. 10.1073/pnas.0506580102.

46. Durinck, S., Spellman, P.T., Birney, E., and Huber, W. (2009). Mapping identifiers for the integration of genomic datasets with the R/ Bioconductor package biomaRt. Nat Protoc 4. 10.1038/nprot.2009.97.

47. Wittig, I., Braun, H.P., and Schägger, H. (2006). Blue native PAGE. Nat Protoc 1. 10.1038/nprot.2006.62.

48. Cogliati, S., Herranz, F., Ruiz-Cabello, J., and Enríquez, J.A. (2021). Digitonin concentration is determinant for mitochondrial supercomplexes analysis by BlueNative page. Biochim Biophys Acta Bioenerg 1862. 10.1016/j.bbabio.2020.148332.

49. Cao, Y., Vergnes, L., Wang, Y.C., Pan, C., Chella Krishnan, K., Moore, T.M., Rosa-Garrido, M., Kimball, T.H., Zhou, Z., Charugundla, S., et al. (2022). Sex differences in heart mitochondria regulate diastolic dysfunction. Nat Commun 13. 10.1038/s41467-022-31544-5.

50. Sánchez-González, C., and Formentini, L. (2021). An optimized protocol for coupling oxygen consumption rates with β-oxidation in isolated mitochondria from mouse soleus. STAR Protoc 2, 100735. 10.1016/j.xpro.2021.100735.

